# Circular extrachromosomal DNA in *Euglena gracilis* under normal and stress conditions

**DOI:** 10.1101/2023.11.29.569251

**Authors:** Natalia Gumińska, Paweł Hałakuc, Bożena Zakryś, Rafał Milanowski

## Abstract

Extrachromosomal circular DNA (eccDNA) enhances genomic plasticity, augmenting its coding and regulatory potential. Advances in high-throughput sequencing have enabled the investigation of these structural variants. Although eccDNAs have been investigated in numerous taxa, they remained understudied in euglenids. Therefore, we examined eccDNAs predicted from Illumina sequencing data of *Euglena gracilis* Z SAG 1224–5/25, grown under optimal photoperiod and exposed to UV irradiation. We identified approximately 1000 unique eccDNA candidates, about 20% of which were shared across conditions. We also observed a significant enrichment of mitochondrially encoded eccDNA in the UV-irradiated sample. Furthermore, we found that the heterogeneity of eccDNA was reduced in UV-exposed samples compared to cells that were grown in optimal conditions. Hence, eccDNA appears to play a role in the response to oxidative stress in Euglena, as it does in other studied organisms. In addition to contributing to the understanding of Euglena genomes, our results contribute to the validation of bioinformatics pipelines on a large, non-model genome.

## Introduction

Extrachromosomal circular DNA molecules of chromosomal origin (eccDNA) are common in Eukarya [1]–[6]. They are well understood in model organisms (including humans), but their presence was confirmed also in other eukaryotes, such as domestic pigeon, sugar beet, and protists (ciliates, trypanosomes) [1], [6]–[12]. Virtually all genomic *loci* can be the source of eccDNAs, although 5’UTRs, exons and CpG islands are pointed out as their hotspots [2], [9]– [11], [13]–[15]. EccDNAs vary in size from a few hundred to a few hundred thousand base pairs and may carry partial or complete genes and/or intergenic sequences [1], [9]–[11], [13], [16]. Some of them are found continuously throughout the cell cycle while others emerge or accumulate under particular conditions [4], [16]–[19].

Circular DNA formation can occur during programmed processes, such as maturation of T-cells and antibody class switching in mammals or regeneration of macronuclei in ciliates [2], [8], [13], [16]. Furthermore, a significant percentage of eccDNA is transposon-derived or linked to the drug resistance acquisition [1], [16], [20], [21]. Thus, the eccDNAs contribute to the plasticity of eukaryotic genomes by increasing its adaptive capacity [2], [22]. However, the accumulation of eccDNA is frequently the result of disease, senescence or stress [4], [9], [13], [14]. In addition, eccDNAs have attracted considerable attention as drivers of oncogenesis often associated with poor prognosis [2], [10], [14], [23]–[25]. They have even been proposed as a diagnostic tool in certain cancers [10], [14], [23], [24].

Recent advances in high-throughput sequencing technologies and computational methods have opened up the possibility to study eccDNA on an unprecedented scale – including non-model organisms lacking the gold-standard genomic data, such as the euglenids [5], [6].

Euglenids, members of Euglenozoa group, are cosmopolitan unicellular algae with potential as a source of biofuels, nutrients, and antioxidants [26]–[29]. *Euglena gracilis* is the best characterized and most frequently used species [30]–[34]. It remains the only representative of the *Euglena* genus with a publicly available genome draft. This preliminary assembly is highly fragmented (N_50_ = 955 bp) and predominantly unannotated [30], [35]. Initially, genomes of euglenids were presumed to be several times larger than the human genome. Currently, the total genome size of *E. gracilis* is estimated to 0.5-2 Gbp [35].

To date, three types of circular DNAs have been identified by molecular methods in *E gracilis*: chloroplast genome, rDNA^circle^ and small mtDNA circles, although the latter remain debatable [36], [31], [37], [38], [32].

Among the Euglenozoans, circular DNA of nuclear origin has been examined only in *Leishmania*, a human pathogens [39]. In these parasites, eccDNAs arise from the chromosomal regions containing inverted repeats [7], [20]. In addition, genomes of *Leishmania* are laden with motifs that evolved from transposable elements and are constantly rearranged [7], [20], [39], [40]. Under selective pressure, the amount of eccDNAs associated with traits beneficial to the cell increases and returns to their original level when the pressure subsides [39]. We hypothesized that similar mechanisms may occur in *Euglena*.

The nuclear-derived circular DNAs of euglenids have not yet been thoroughly studied. To address this issue, we examined eccDNA candidates predicted from the short-read sequencing data of eccDNA-enriched libraries isolated from *E. gracilis* cells cultured under optimal growth conditions and periodically exposed to UV irradiation. Our work contributes to a deeper understanding of the organization and plasticity of genomes of *Euglena*. Finally, it validates bioinformatics tools on a relatively large, complex, non-model genome.

## Materials and methods

### Cell strains and growth conditions

Cultures of *Euglena gracilis* Z strain (SAG 1224–5/25) from the collection of Institute of Evolutionary Biology (University of Warsaw, Poland) were grown statically in a transparent glass tubes with a liquid Cramer-Myers medium as described elsewhere [41] until early logarithmic growth phase.

### UV irradiation

To examine the effect of UV light exposure on eccDNA formation in *E. gracilis*, an initial liquid cultures (10^5^ – 10^6^ cells ml^-1^) were diluted to a density of 1.6 × 10^4^ cells ml^-1^, and 200 μl of the cell suspensions were inoculated into 3 ml of Cramer-Myers medium in transparent glass tubes. These cells were then irradiated with UV-A and UV-B for one hour a day for 5 consecutive days [42] in UV crosslinker (cat. no: 89131-486, VWR). On each exposure, cells were applied a radiation dose of 0.36 J cm^-2^. All other light was obscured during the irradiation periods.

Cell morphology was assessed using Nikon Eclipse E-600 light microscope with differential interference contrast, equipped with the NIS Elements Br 3.1 (Nikon, Japan) software.

### Total DNA isolation

For each condition, 1 × 10^10^ cells were pelleted (5 min, 3,000 rpm). Total DNA was isolated from *E. gracilis* cells using cetyltrimethylammonium bromide (CTAB; AppliChem) according to the protocol optimized for the euglenids as described previously [41]. The total DNA extraction was performed before the enrichment to remove the secondary metabolites [43], which could affect the recovery of eccDNA fraction [41]. The integrity of the DNA was assessed with electrophoresis in a 1.5% agarose gel stained with 0.5% Midori Green (Nippon), run in 1× TAE buffer and visualized with the ChemiDoc UV transilluminator (Bio-Rad).

### EccDNA enrichment

Purification of eccDNA from *E. gracilis* was optimized based on the Circle-Seq eccDNA method for budding yeast [16]. In brief, 50 μg of total DNA in 10 replicates of 5 μg each per experimental condition were used for circular DNA enrichment with GeneMATRIX Plasmid Miniprep DNA Purification Kit (EURx) according to the manufacturer’s instructions. DNA was eluted with 50 μl of elution buffer heated to 80°C. According to the High Sensitivity dsDNA Assay by Qubit 3.0 fluorometer (Thermo Scientific), the 236–449 ng of eccDNA enriched DNA were obtained.

Remaining linear DNA molecules were digested with Plasmid-Safe ATP-dependent exonuclease (Epicentre) continuously at 37°C for 120 h. To ensure complete hydrolysis of linear DNA, the reaction mixture was supplemented with an additional ATP and DNase every 24 h (16 units per day) according to the manufacturer’s protocol. Finally, the enzyme was heat inactivated for 30 min at 70°C.

Using a 2100 Bioanalyzer (Agilent) it was estimated that the initial amount of total DNA taken for the analysis has decreased by 4.5 × 10^5^-fold in untreated and 6 × 10^6^-fold in an UV-irradiated sample. Which, given the differences in genome size, is consistent with data obtained in other studies for yeast and human cells [16], [44].

Complete removal of linear DNA was confirmed by PCR of the *tubA* gene. Amplification with rDNA^circle^ complementary primers confirmed their presence in the samples.

### Polymerase chain reactions

The *tubA* genes were used to test whether the linear DNA was digested completely. The rDNA^circle^ was used to check whether the circular DNA remained intact. Briefly, 2 μl of samples after exonuclease digestion were used as a template in the reaction prepared as described elsewhere [34], [45].

The following primers were used for *tubA* amplification:

TAe2F – AGCGCCCCACTTACACCAACCTGAAC

TAi6R – GGGGGTCGGATCGGGTGCTATG

The following primers were used for rDNA^circle^ amplification:

LSU-F – CCCCGATTCATGCACCAAGTCT

LSU-R – CGCACTGGGAGACGATACACTTCA

### Library preparation and sequencing

The eccDNA isolates were pulled (each condition separately), precipitated and amplified using Illustra TempliPhi kit (GE Healthcare) according to the manufacturer’s instructions. Fragmented genomic DNA libraries were produced from 10 ng of each sample treated with 1.5 µl of Nextera XT tagmentase (Illumina) at 37°C for 45 min with gentle shaking. The DNA libraries were size-selected on AMPure beads (NEB) and assessed using a 2100 Bioanalyzer (Agilent), followed by sequencing on the HiSeq 4000 (Illumina) platform in PE150 mode. Library preparation, quality control, and sequencing were outsourced (Genomed, Warsaw, Poland).

### Preprocessing and quality assessment of reads

Raw sequencing reads were quality-checked using the FastQC 0.11.9 software [46] and preprocessed (trimmed, quality filtered) with Atria 4.0.0 [47]. Trimmed reads were mapped to the reference draft genome with bwa-mem2 with default parameters [48]. Resulting sequence alignment, after conversion to fasta format, was used for BLAST search against the NCBI GenBank database (blastn, -outfmt “6 qseqid staxids bitscore std” -max_target_seqs 1 - max_hsps 1 -num_threads 4 -evalue 1e-25). The resulting data was used to determine the origin of the reads using Blobtools [49].

### Comparison of eccDNA detection pipelines

In all cases the draft genome assembly of *Euglena gracilis* retrieved from the European Nucleotide Archive (ERP109500) was used as reference for the genome-guided eccDNA screening [35]. The following software was tested: Circle-Map (https://github.com/iprada/Circle-Map), ecc_finder (https://github.com/njaupan/ecc_finder) and ECCsplorer (https://github.com/crimBubble/ECCsplorer/). The Circle-Map was launched in both Realign and Repeats modes, with the default parameters [3], [4]. The ecc_finder was launched in a mapping mode guided by a reference genome and in reference-free, *de novo* assembly mode [5]. In reference-dependent mode, the minimap2 was applied instead of the bwa to speed up the computation (--aligner minimap2) with otherwise default parameters [5], [50]. The ECCsplorer was launched with the trimming option disabled, read count set to auto (with a genome coverage 0.1×), an estimated genome size of 500 Mbp. Both guided and *de novo* modes were applied. As control for the reference-free eccDNA detection, whole-genome sequencing reads from the same strain, used for assembly of the genome draft were used (ERR2660554).

The data in BED format were retrieved from the outputs of the Circle-Map and ecc_finder, and compared using BEDtools intersect (-u, -f 0.90, -r options)[51]. The results were visualized using the ggplot2 [52] R package.

### Mappping to selected *E. gracilis* sequences

Reads corresponding to the chloroplast genome of *E. gracilis* were mapped with NOVOPlasty software using NC_001603.2 sequence retrieved from NCBI GenBank both as guide and seed [53].

The rDNA^circle^ reference sequence was curated manually based on the X53361.2 sequence retrieved from NCBI GenBank and reads from whole-genome sequencing performed as part of another project [41]. Trimmed reads were then mapped to the resulting sequence with bwa-mem2 using default parameters [48].

The set of 79 mitochondrial contigs of *E. gracilis* was curated based on the publicly available draft assembly and reads obtained as part of another project [41] (https://doi.org/10.18150/THCE6Y). Trimmed reads were mapped to the resulting sequence with bwa-mem2 using default parameters [48].

### Hardware and software

Computations were done on Unix machine (Ubuntu 20.04.3 LTS, Intel Xeon Platinum 8268 ×64 CPU 2.90 GHz, 96 cores, 1008 GB RAM) with Python 3.7.9, Biopython [54], Scipy [55], pyRserve [56], R 4.1.2 with following packages: Bioconductor 3.14, dplyr [57], ggplot2 [52], gridExtra [58], ggrepel [59] and other third-party software: BEDtools [51], Blast+ [60], Blobtools [49], bwa-mem2 [48], IGV [61], minimap2 [50], RepeatExplorer2 [62], samtools [63], segemehl [64], seqtk [65], Tablet [66].

### Availability of data and resources

The data presented in the study were deposited in European Nucleotide Archive (ENA) repository under project number PRJEB67978 (ERP152989). All propagative materials (e.g. euglena cultures) that were used to obtain results presented in the article will be available on request.

## Results

### EccDNA enrichment protocol for euglenids

First, we established a protocol for the eccDNA isolation from euglenids. To accommodate the peculiarity of *Euglena*, we modified the eccDNA fractionation protocol of Møller et al. [16], [44]. We had to account for the unique cell cover, the accumulation of secondary metabolites, and the genome size (Fig. 1). Thus, we isolated total DNA from *E. gracilis* using our CTAB-based method, which has already been proven effective for high-throughput applications [41]. Then, we performed two alkaline lyses: before and after digestion of linear molecules with ATP-dependent exonuclease. Thereby, we wanted to increase the probability of preparing the libraries exclusively from circular molecules. Furthermore, by performing a second minilysis instead of applying rare-cutting endonucleases, we prevented the quality of the samples from being compromised (Fig. 1A). According to our experience, algal genetic material is particularly susceptible to degradation if subjected to multiple procedures [41], [67]–[69].

**Figure 1.**
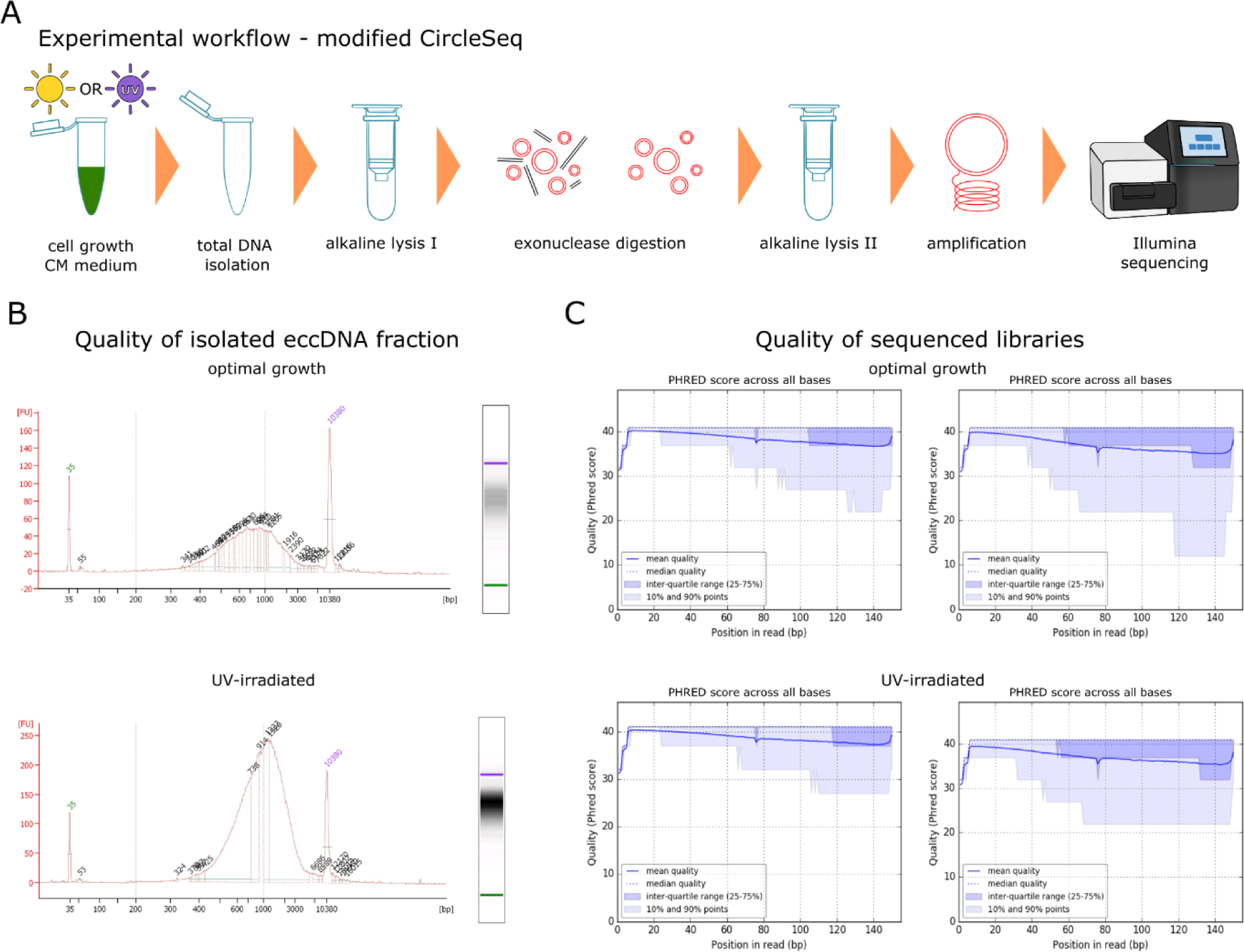
Library preparation and quality control. (A) Overview of the experimental fractionation of eccDNA with modified CircleSeq protocol. Extra steps were introduced to accommodate unique morfological features of *Euglena* cells. (B) Results of concentration and quality assessment of eccDNA-enriched samples with 2100 Bioanalyzer prior sequencing. (C) PHRED quality score values (Illumina 1.9 encoding) across all bases for 150 PE HiSeq Illumina reads at each position in the sequnced libraries.

Final DNA concentrations in both samples were consistent with values reported for eccDNA-fractionated samples from other organisms [2], [12], [18]. The majority of the molecules in our samples ranged in size from 600 to 1,000 bp, with slightly larger eccDNA candidates predominating in the UV-exposed one. In addition, the latter contained approximately 3 times as much potentially circular molecules (Fig. 2A).

**Figure 2.**
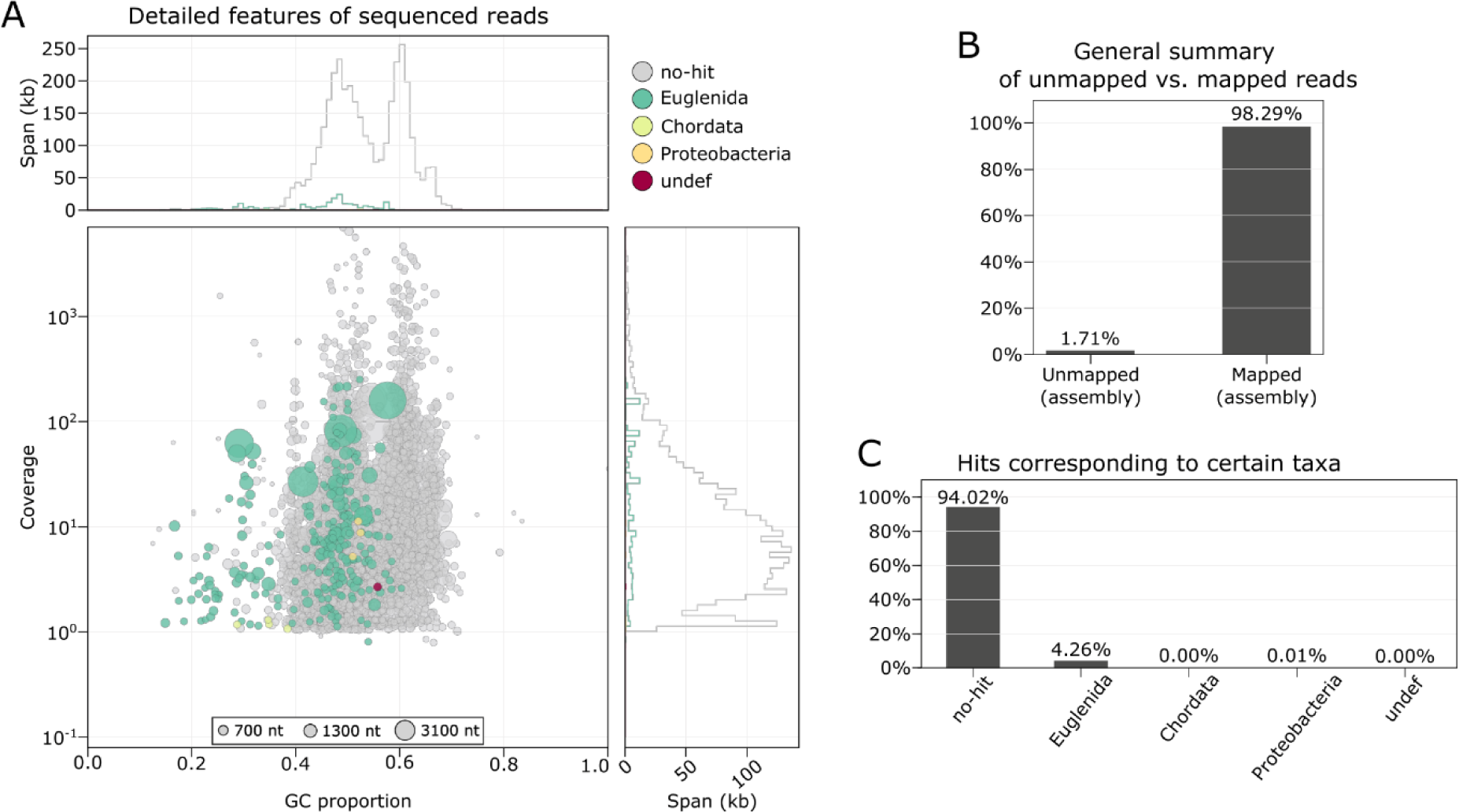
Visualisations of combined assembly of both sequenced libraries. (A) BlobPlot of the reads mapped to the draft genome of *E. gracilis* and queried against NCBI nucleotide database. Circles illustrating assembly sequences are scaled colored according to the sequence length and coloured by taxonomic annotation based on blastn search. The circles are positioned on the x-axis by their GC-content and on the y-axis by to the overall coverage in both libraries. (B) ReadCovPlot showing the proportion of either unmapped or mapped reads. (C) ReadCovPlot showing the percentage of mapped reads by taxonomic group.

The sequencing library corresponding to optimal growth conditions produced more than 230 million reads, while the UV-irradiated cells produced nearly 220 million reads (Table 1). Both runs were of high quality, as verified by FastQC: the mean PHRED scores exceeded 35, with the majority of nucleotides within the reads reaching the highest scores (Fig. 1C).

**Table 1.** Basic parameters of sequencing results.

|  | **optimal growth** |  | **UV-irradiated** |  |
| --- | --- | --- | --- | --- |
| **File name** | EGZ_steady_L1 | EGZ_steady_L2 | EGZ_UV_L1 | EGZ_UV_L2 |
| **Number of reads** | 231 604 923 | 231 604 923 | 219 852 679 | 219 852 679 |
| **Number of bases** | 31 639 830 926 | 29 495 372 117 | 30 523 226 074 | 28 633 445 168 |
| **Read length** | 15-150 | 15-150 | 15-150 | 15-150 |
| **% GC** | 55 | 55 | 54 | 53 |

### Taxonomic assignment of sequencing data

We then checked what percentage of sequencing reads actually derived from *E. gracilis*. We mapped the sequencing reads to the draft genome and screened for potential contaminants (Fig. 2). The resulting alignment was highly fragmented, consisting mostly of short contigs (< 700 bp). Such high dispersion is a common feature of eccDNA-enriched libraries, suggesting the presence of discordant and split-end reads [2], [12], [16], [44]. For most contigs, we achieved reasonably good coverage given the size of the *E. gracilis* genome (Fig. 2A). A majority of the mapped fragments contained 40-60% GC nucleotides, with two distinct subpopulations visible. In addition, a third less abundant cluster with significantly lower GC content could also be noted. It most likely corresponds to chloroplast and mitochondrial fractions.

Remarkably, more than 98% of reads were mapped, indicating high purity of the library (Fig. 2B). Striking majority of the contigs could not be taxonomically assigned with a BLAST search with only about 4% assigned to euglenids (Fig. 2C). This is due to the poor representation of *Euglena* (including *E. gracilis*) in the GenBank database and incomplete genome annotation.

### Observation of known circular DNAs in *E. gracilis*

Small amounts of chloroplast reads were detected in the sequenced samples. However, they were not sufficient to assemble the entire chloroplast genome. Since the cpDNA has a size of 143,170 bp, it can be assumed that the chloroplast DNA was fragmented during preparation and/or was too large to be successfully isolated and included in the library.

A substantial fraction of the libraries (∼1% under optimal conditions and ∼3% after UV-irradiation) corresponded to the rDNA^circle^. In both libraries, the reference sequence (11,055 bp) was fully covered (Fig. 3). Remarkably, inspection of the assembly graphs suggests heteroplasmy of the rDNA circles, consistent with our previous studies [70].

**Figure 3.**
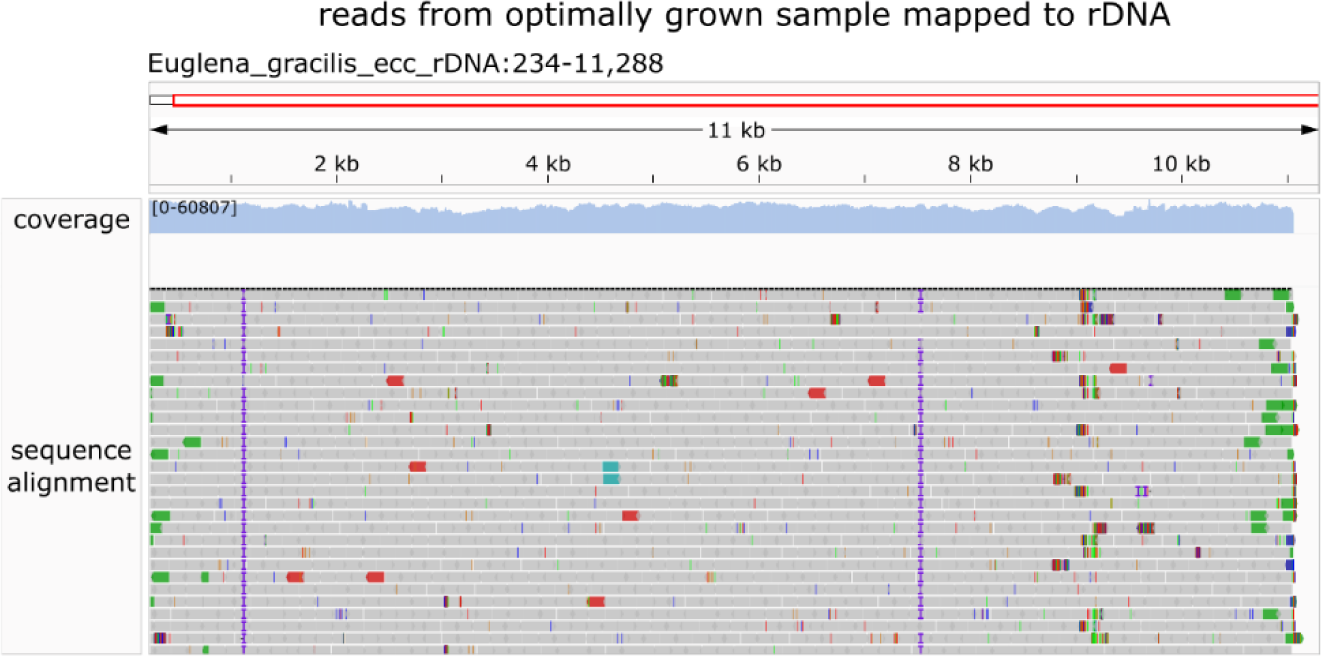
Integrative Genomics Viewer (IGV) snapshot of sequence alignment of reads mapping to the manually curated reference sequence of rDNA^circle^. Split-end reads in green, discordant reeds in red. Library was prepared from cells cultured in optimal conditions.

### Benchmarking of eccDNA detection pipelines with *E. gracilis* data

EccDNA candidates are identified based on read properties revealed by the mapping to the reference genome, such as high coverage and the presence of split-end and discordant reads [3]–[6]. The reference-free approaches also had been proposed [5], [6], [71]. We selected three tools: Circle-Map [16], ecc_finder [5], and ECCsplorer [6] as the most promising to handle our data. These algorithms have been previously applied to numerous eukaryotes, including non-model organisms with large genome sizes and poorly annotated or fragmented assemblies [3]– [6], [11], [16].

Circle-Map run in Realign mode (identifying putative circular DNA junctions) failed to identify eccDNA candidates in any of the samples. However, in Repeats mode (identifying putative repetitive circular DNA) it indicated around 2 thousand eccDNA candidates per sample. Specifically, 2274 potential DNA circles were identified in optimal growth conditions, and 1865 under the UV-irradiation (Table 2). All of these putative circles were supported by at least 10 repeated reads. After filtering the results based on coverage increase and coverage continuity, as suggested by Møller et al. [4], [11], [16], we obtained 872 circles corresponding to optimal conditions and 612 corresponding to UV-irradiation (Supplementary table 1). Whereas ecc_finder run in genome guided mode reported 698 circles in optimally cultured and 639 circles in UV-exposed *E. gracilis* (Table 2). Of these, 325 and 288 candidates, respectively, were represented by at least 10 discordant reads (Supplementary table 2).

**Table 2.** Comparison of predictions provided by applied eccDNA detection tools. The ‘-‘ indicate runs finished with errors preventing from obtaining final results. The table contains raw predictions.

| **pipeline** | **mode** | **putative eccDNAs**<br>**optimal growth** | **putative eccDNAs**<br>**UV-irradiated** |
| --- | --- | --- | --- |
| Circle-Map | Realign | 0 | 0 |
| Circle-Map | Repeats | 2274 | 1865 |
| ecc_finder | genome guided | 698 | 639 |
| ecc_finder | reference free | - | - |
| ECCsplorer | genome guided | - | - |
| ECCsplorer | reference free | - | - |

Unfortunately, ecc_finder was unable to complete the run in the genome-unassisted mode. The analyses with ECCsplorer also failed to complete (Table 2). Therefore, we focused on the results produced by Circle-Map and ecc_finder.

### Characterization of eccDNA candidates in *E. gracilis*

We then decided to characterize the predicted eccDNA candidates as best as possible based on the available draft genome of *E. gracilis*. The putative eccDNAs predicted by Circle-Map are about twice as long as the circles predicted by ecc_finder in both culture conditions (Fig. 4A). The results from each software considered separately are consistent in both libraries. The divergences between the outputs of the used tools are due to the different algorithmic approaches underlying these pipelines.

**Figure 4.**
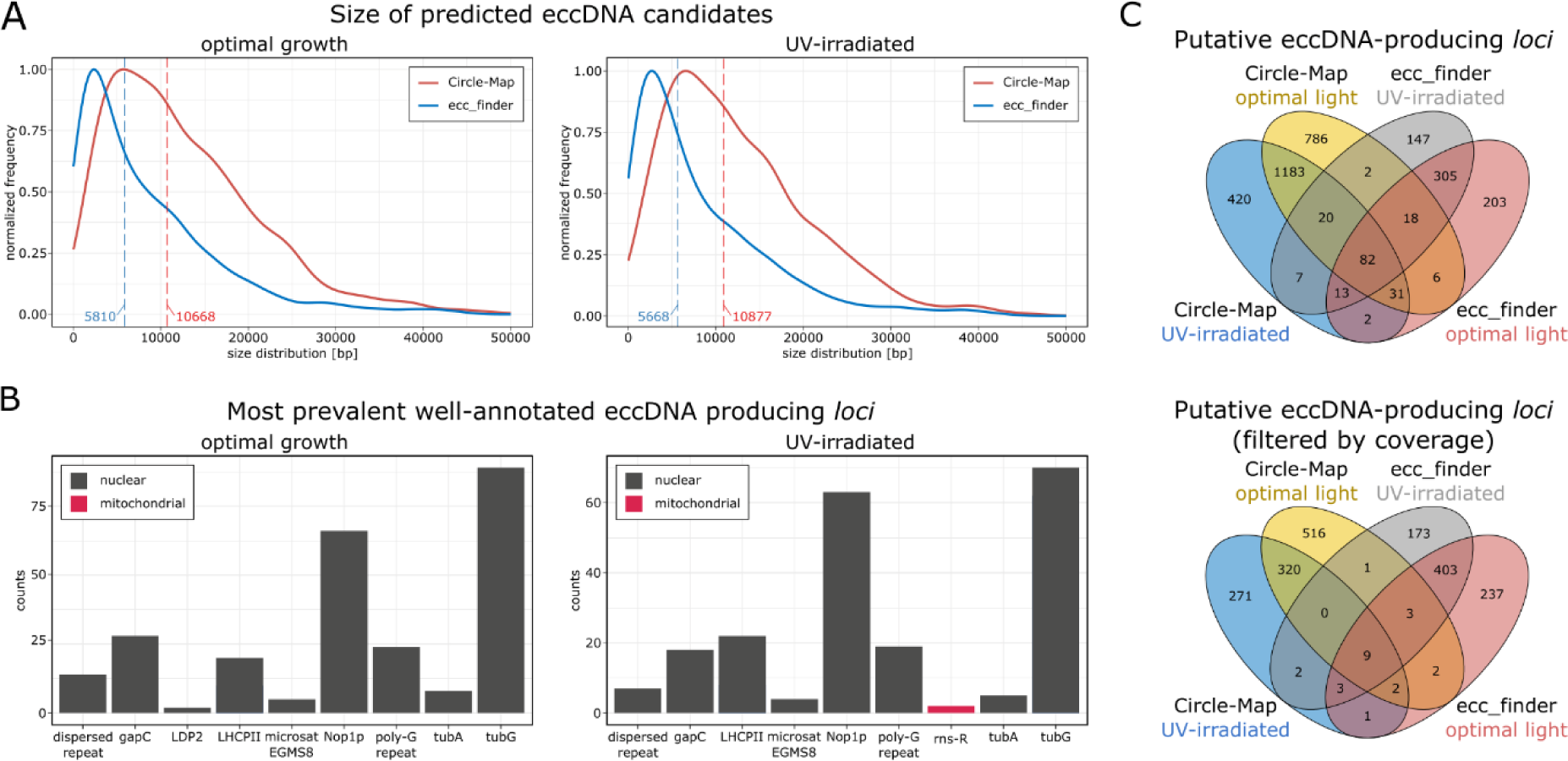
Characteristics of predicted eccDNA candidates. (A) EccDNA length distribution under normal and UV-stress conditions. Predictions made by each pipeline are plotted separately. Median eccDNA lengths are shown as vertical dashed lines. (B) Most prevalent sources of putative eccDNA in draft genome of *E. gracilis*. Experimental conditions are plotted separately. (C) Venn diagrams comparing amounts of potential eccDNA-producing *loci*. Upper diagram shows unfiltered data, while the bottom diagram shows predictions filtered based on coverage increase and coverage continuity.

**Figure 5.**
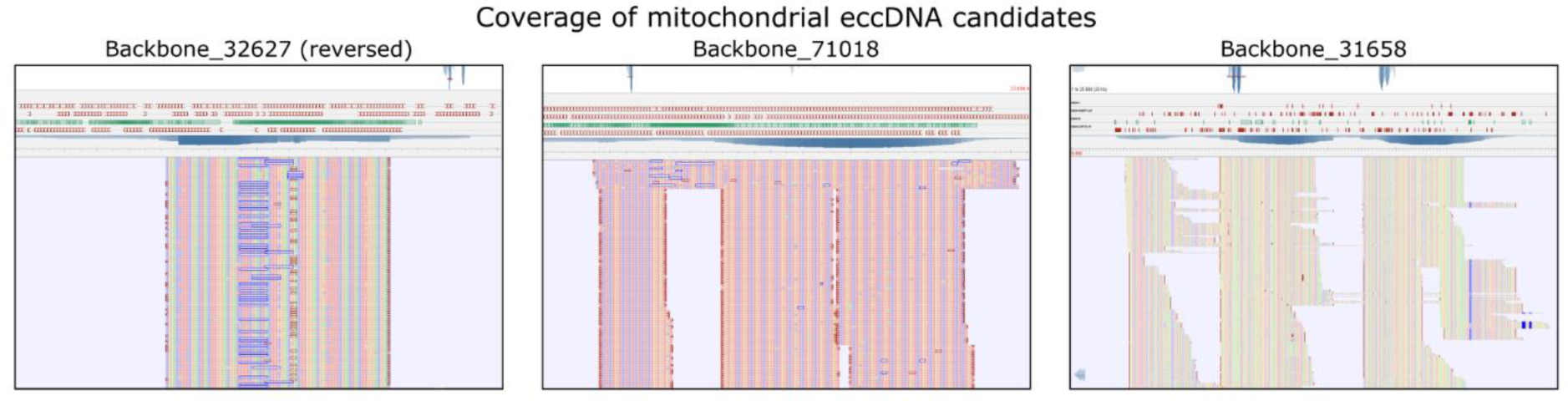
Tablet snapshots of sequence alignments of reads mapping to the manually curated reference mitochondrial contigs. Repetitive regions seem to be the hot-spots for eccDNA formation in mitochondrial DNA. Soft-clipping reads (split) are visible. The UV-irradiated sample was shown as an example.

Since most backbones in the reference assembly are unannotated, we BLAST-searched the results against the NCBI Genbank database. In this way, we identified repetitive regions, microsatellites and several housekeeping genes (e.g. tubulins, fibrilarin) as potential sources of circular eccDNAs in the nuclear genome of *E. gracilis* (Fig. 4B). Sequence alignments indicate that a significant percentage of these DNA circles contain intron sequences and/or their flanks.

In the next step we identified potential hotspots of eccDNA formation. We focused on genomic *loci*, as multiple potential circles could originate from the same site due to the presence of repetitive sequences.

For a given tool, the sets of eccDNA candidates from the two experimental conditions are largely consistent with each other (Fig. 4C). Even after filtering the results based on coverage increase and continuity, about 20% of the putative circles are common for optimal growth and UV exposure. However, only a small portion of the results are common to both tools. This is particularly evident after filtering out the data (Fig. 4C).

### EccDNA candidates in mitochondrial genome of *E. gracilis*

To our surprise, mitochondrial DNA was also among the potential sources of eccDNA in the UV-irradiated sample (Fig. 4B). Especially since the literature data on the conformation of mitochondrial DNA of euglenids are contradictory [32], [37], [38].

We investigated this further using our revised set of 79 mitochondrial contigs of *E. gracilis*. Thus, we identified 20 loci potentially giving rise to eccDNA. Of these, 180 eccDNA candidates derived from optimal culture conditions and 168 from the sample after UV exposure. Potential mitochondrial circles, after being mapped to our revised mitochondrial contigs, had coverages an order of magnitude higher than nuclear circles mapped to the publicly available reference draft assembly (Supplementary table 3).

## Discussion

Exploring the genomes of euglenids is challenging due to their considerable size, complexity, and the difficulty of obtaining high-quality material for sequencing [41]. Here we present the first protocol for eccDNA fractionation optimised for *E. gracilis*, which not only paves the way for further studies on euglenids, but also on other algae.

Although the draft genome of *E. gracilis* is largely uncurated, the eccDNA repertoire of this species can be studied on a fairly high extent. However, in presenting preliminary eccDNA screening results, we are aware of the limitations inherent in detailed analysis without a high-quality reference genome.

Our study highlights the necessity for a customized computational approach for the eccDNA discovery in euglenids. It is important to consider the limitations of pipelines intended for reference-free eccDNA reconstruction. In particular, given that the similarities between the eccDNA candidates and repetitive regions, structural variants and assembly errors may render current methods prone to false positives [5], [6], [22]

Our findings show dynamic changes in the eccDNA repertoire of *E. gracilis* cells grown under optimal light exposure and under periodic UV irradiation. We observed a limited number of shared DNA circles between the experimental conditions, suggesting that eccDNA formation is involved in the genomic response to external factors. As a result of UV irradiation, the heterogeneity of eccDNA decreased significantly, which is consistent with findings in other organisms [4], [17]. This also suggests that the eccDNA landscape responds to environmental stressors with a selective retention of certain eccDNA molecules [2], [4], [12].

Remarkably, our study provides initial insights into potential mitochondrial circular molecules in *E. gracilis*, challenging existing views on the mitochondrial genome’s structure (https://doi.org/10.1093/gbe/evv229). However, detailed analysis remains contingent on a comprehensive and revised annotation.

In conclusion, this research lays a solid foundation for future studies on structural variants and DNA circles in *E. gracilis*. It not only introduces a novel protocol for eccDNA fractionation but also underscores the urgency of refining the reference genome to unlock the full potential of genomic studies in this unique organism.

## Funding

This work was supported by the 2017/27/N/NZ2/01321 grant from Polish National Science Centre to NG.

## Author contributions

NG and RM contributed to the conception and design of the study. BZ provided and maintained the cell cultures. PH and RM curated the reference assembly of *E. gracilis*. NG conducted the experiments, performed bioinformatical analysis, visualized the data and wrote the first draft of the manuscript. RM, PH and BZ contributed to manuscript revision. All authors read and approved the submitted version.

## Conflict of interest

The authors declare that the research was conducted in the absence of any circumstances that could lead to a potential conflict of interest.

## Acknowledgements

The authors thank prof. Andrzej Dziembowski for proofreading and discussion and Mariusz Czarnocki-Cieciura for the hardware resources.

## Supplementary data

**Supplementary table 1**

Merged output of Circle-Map pipeline (for both experimental conditions), filtered by coverage increase and continuity. Draft genome assembly was used as reference.

**Supplementary table 2**

Merged output of ecc_finder pipeline (for both experimental conditions). Draft genome assembly was used as reference.

**Supplementary table 3**

Merged output of Circle-Map pipeline (for both experimental conditions). Manually curated set of mitochondrial contigs was used as reference.

## Notes

https://www.ebi.ac.uk/ena/browser/view/PRJEB67978

### Competing Interest Statement

The authors have declared no competing interest.

